# Distinct learning, retention, and generalization in de novo learning

**DOI:** 10.1101/2023.10.02.560506

**Authors:** Raphael Q. Gastrock, Bernard Marius ’t Hart, Denise Y. P. Henriques

## Abstract

People correct for movement errors when acquiring new motor skills (de novo learning) or adapting well-known movements (motor adaptation). These two motor learning types should be distinct, as de novo learning establishes new control policies while adaptation modifies existing ones. Here, we distinguish between these two motor learning types, and assess de novo learning retention and generalization. In study 1, participants train with both 30° visuomotor rotation and mirror reversal perturbations, to compare adaptation and de novo learning respectively. We find no perturbation order effects, and that learning develops with similar rates and comparable asymptotes for both perturbations. Explicit instructions also provide an advantage during early learning in both perturbations. However, mirror reversal learning shows larger inter-participant variability. Furthermore, movement initiation is slower for the mirror perturbation, and we only observe reach aftereffects following rotation training. In study 2, we use a browser-based mirror reversal task to investigate learning retention and generalization to the untrained hand and across the workspace. Learning persists across three or more days, substantially transfers to the untrained hand, and to targets on both sides of the mirror axis. Our results show that behavioral mechanisms underlying motor skill acquisition are distinct from adapting well-known movements.

## Introduction

Moving appropriately requires that people learn from movement errors. Such error-processing is often classified into two motor learning types. One is motor adaptation, where we modify an existing control policy to regain previous levels of performance ^[1–4]^. The other is skill acquisition, or de novo learning, where we must establish a new control policy ^[4–6]^. Previous studies have compared both motor learning types ^[5,7–11]^, but there are still several aspects of de novo learning that warrant investigation, including its retention and generalization. Here, we investigate adaptation and de novo learning using visuomotor rotation and mirror reversal perturbations respectively. Further, we investigate retention and generalization of learning in a browser-based mirror reversal task. To better capture the differences between these two motor learning types, we use a within-subjects design and a large sample size in two studies. Both experiments extend our understanding of how error-processing in motor adaptation is distinct from de novo learning.

In reaching movements, errors introduced from visual ^[1,12–14]^ or mechanical perturbations ^[15–18]^, are gradually adapted in subsequent trials. These adaptive changes involve updating internal forward models based on sensory prediction errors, which are actual sensory consequences compared with predictions from the efference copy of the outgoing motor command ^[2,4,19–20]^. This remapping is manifested through reach aftereffects or persistent hand movement deviations after perturbation removal ^[2,4,21]^. While aftereffects are considered evidence of implicit adaptation ^[1,12]^, explicit processes also account for these adaptive changes and are expressed as cognitive strategies to compensate for the perturbation ^[6,11,22–26]^. Previous studies have investigated de novo learning using a mirror reversal perturbation, where cursor visual feedback is in the flipped direction of the hand movement relative to a mirror axis ^[4–8,10–11,27–30]^. For this perturbation, sensory prediction error-based implicit adaptation is counter-productive and leads to larger errors, thereby requiring more time to establish a new movement control policy ^[5,29,31–34]^. Consequently, reach aftereffects are not typically observed ^[5,9,29]^, as one can simply switch between control policies upon perturbation removal. Explicit processes, therefore, contribute greatly to de novo learning ^[6–8,10–11,35]^, and manifest as slower movements early in de novo learning as compared to adaptation ^[5]^. These learning characteristics distinguish the underlying mechanisms between motor adaptation and de novo learning.

As adaptation and de novo learning progress differently, learning retention is also distinct. Adaptation quickly decays over time ^[36]^, and is only partially retained ^[4,37–38]^. Aftereffects also wash out and revert to baseline levels of reaching within a few trials ^[4,39]^. Savings, or faster learning rates when re-experiencing the perturbation, show how people still commit large initial errors upon experiencing the perturbation a second time ^[40–41]^. In de novo learning, retention of learning persists across a few days for reaching tasks ^[5,10–11]^, and up to a month for continuous target tracking tasks ^[6–7]^. Furthermore, no aftereffects are expected following de novo learning, such that no de-adaptation occurs ^[5,9,29]^. Instead of savings, offline gains are expected, where initial performance during the second session starts off at the same or even better level as asymptotic learning in the first session ^[5]^. Thus, de novo learning seems to be more persistent than adaptive changes.

Another characteristic of learning is how well it transfers to different conditions. In adaptation, aftereffects show narrow generalization patterns that peak towards the trained movement direction ^[1,4,42–43]^. Intermanual transfer, where less initial errors or faster learning are observed during training with the untrained hand, is usually asymmetric or transfers from the dominant to non-dominant hand but not vice-versa ^[44–48]^. In de novo learning, implicit contributions to learning also seem to be specific to the trained movement direction ^[10,29]^. Intermanual transfer with reversed feedback of the actual hand, however, seems to not be asymmetric and even facilitates transfer of learning to the untrained hand ^[49–50]^. Here, we test whether learned performance is specific to target location relative to the mirror axis, and whether this learning transfers across the workspace and to the untrained hand. Understanding the generalization patterns across movement directions and effectors in a de novo learning task distinguishes it further from motor adaptation.

We confirm distinctions between adaptation and de novo learning in two studies. In study 1 (*N* = 32), participants train with both a 30° visuomotor rotation and mirror reversal perturbation (Fig. 1A-1D). Additionally, half of the participants received instructions about the nature of each perturbation and a strategy to counter for it. We hypothesize that learning each perturbation does not affect the other, that different behavioral measures are distinct between the two perturbations, and that explicit strategies provide an immediate advantage in learning for both perturbations. For study 2 (*N* = 63), we collect data from large sample sizes in a browser-based experiment, to investigate the retention and generalization of de novo learning. In session 1, participants train with targets in the upper-right quadrant of the workspace (quadrant 1; Fig. 1E-1F), that have different distances relative to a vertical midline mirror axis. Participants then return after a minimum of three days, and we test them on the same target locations, before testing on corresponding targets within the lower-right and upper-left quadrants of the workspace, followed by reaches using their opposite untrained hand. We hypothesize that retention and generalization patterns in the mirror reversal task will be distinct from patterns previously observed in adaptation.

**Figure 1.**
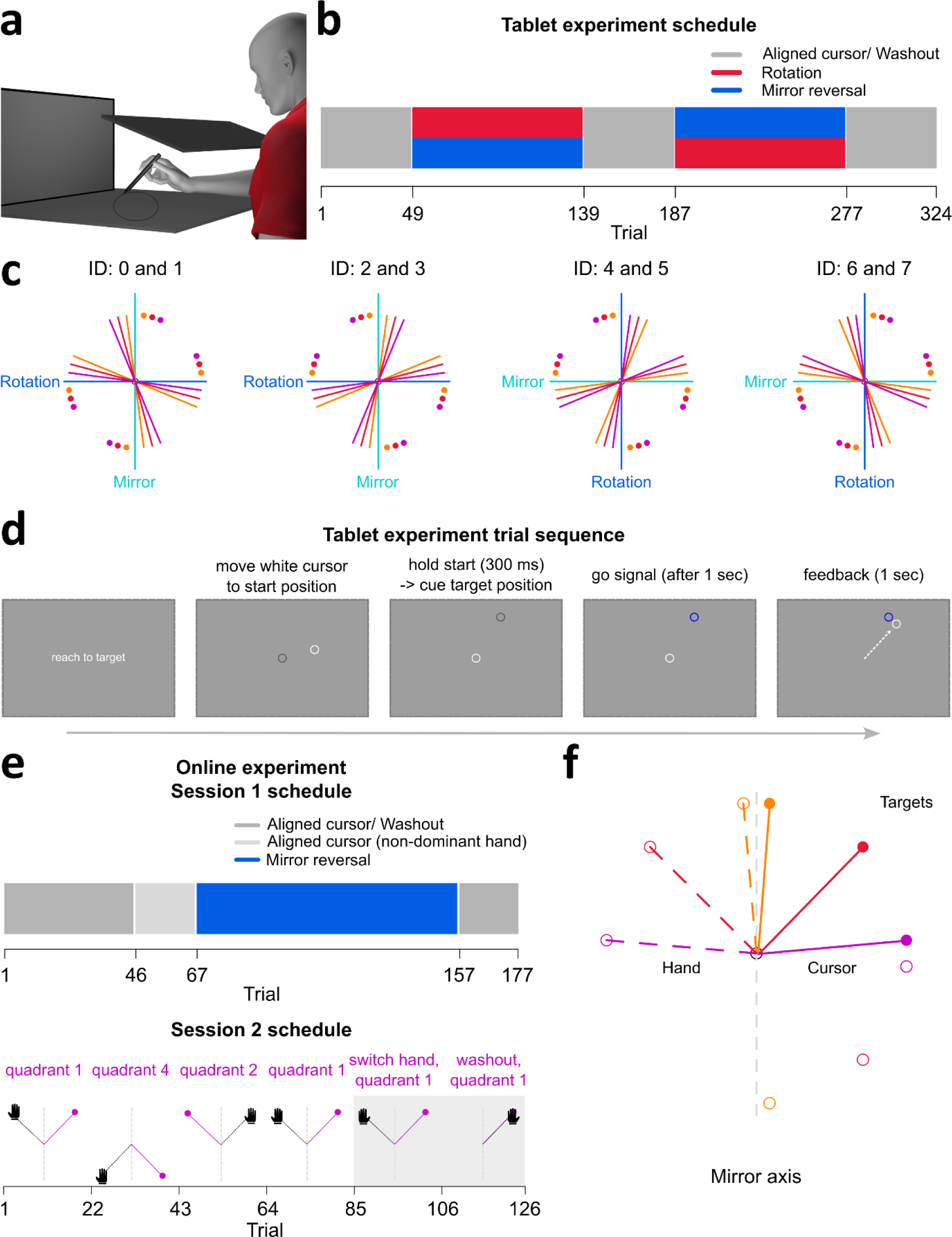
Tablet and online experimental set-up. (**a**) Participants used a stylus to move across a digitizing tablet, while a monitor displayed stimuli and visual feedback of their hidden hand position. (**b**) Aligned cursor, or baseline, reaches had matched cursor and hand positions. Participants (N = 32) completed 48 aligned cursor trials, followed by 90 training trials with either rotation or mirror reversal perturbations, and 48 washout trials. They then completed 90 training trials with the other perturbation, followed by 48 washout trials. (**c**) Even numbered participant IDs experienced the rotation before the mirror reversal, and vice-versa for odd IDs. Each perturbation type corresponded to either the vertical or horizontal midline axis and was counterbalanced across participants. Each axis had six target locations, either 7.5°, 15°, or 22.5° away from both ends of the axis on its positive or negative side. Colored dots indicate targets, and solid lines show the corresponding correct hand movement directions. (**d**) Participants moved the stylus to bring a white cursor to the centre start position. They waited for a go signal (target turned blue) before moving to the cued target location. The cursor remained visible throughout the reach, and they were instructed to hold their terminal position. (**e**) In session 1 (N = 63), participants completed 45 aligned cursor reaches with their dominant hand, before switching to 21 aligned reaches with their non-dominant hand. With their dominant hand, they completed 90 mirror reversed training trials, followed by 21 washout trials. In session 2 (N = 48), participants immediately performed 21 mirror reversed trials with their dominant hand to quadrant 1 targets. Target locations were then switched to quadrant 4, followed by quadrant 2, before performing top-up reaches in quadrant 1 (21 trials/ quadrant). They switched to their non-dominant hand to perform 21 mirror reversed trials to quadrant 1 targets, followed by 21 washout trials. (**f**) Display of target locations, either 5°, 45°, or 85° (near, middle, far targets) away from the vertical mirror axis. Colored dashed lines indicate hand movement direction, and corresponding solid lines indicate cursor direction. Filled dots show quadrant 1 targets, unfilled dots indicate quadrant 2 and 4 targets.

## Results

### Distinct learning between adaptation and de novo learning

In the tablet experiment, we implement a within-subjects design where participants encounter both rotation and mirror reversal perturbations. We therefore test for a perturbation order effect to confirm the independence of learning in each perturbation. To do so, we use an exponential decay function (details in Methods) that estimates the rate of change and asymptote of learning in each perturbation. Using paired t-tests that compare these parameters, we find no evidence for a perturbation order effect for either the rate of change, nor asymptote, for both the rotation (rate of change: BF = 0.486; asymptote: BF = 0.429) and mirror reversal (rate of change: BF = 0.606; asymptote: BF = 0.835; detailed statistics in R notebook ^[51]^). These results suggest that learning between the two perturbation types is independent.

As learning is independent in the two perturbation types, we compare learning rates and asymptotes between the rotation and mirror reversal perturbations. Participants learn both perturbation types within 90 trials, but learning in the mirror reversal has larger inter-participant variability (Fig. 2A). Using a paired t-test to compare exponential decay function parameters between the two perturbation types, we find no evidence that the two perturbations differ in both parameters (rate of change: BF = 0.365; asymptotic learning: BF = 1.836). Thus, while learning for the mirror reversal is much more variable, it develops equally quickly and to a comparable asymptote as rotation adaptation.

**Figure 2.**
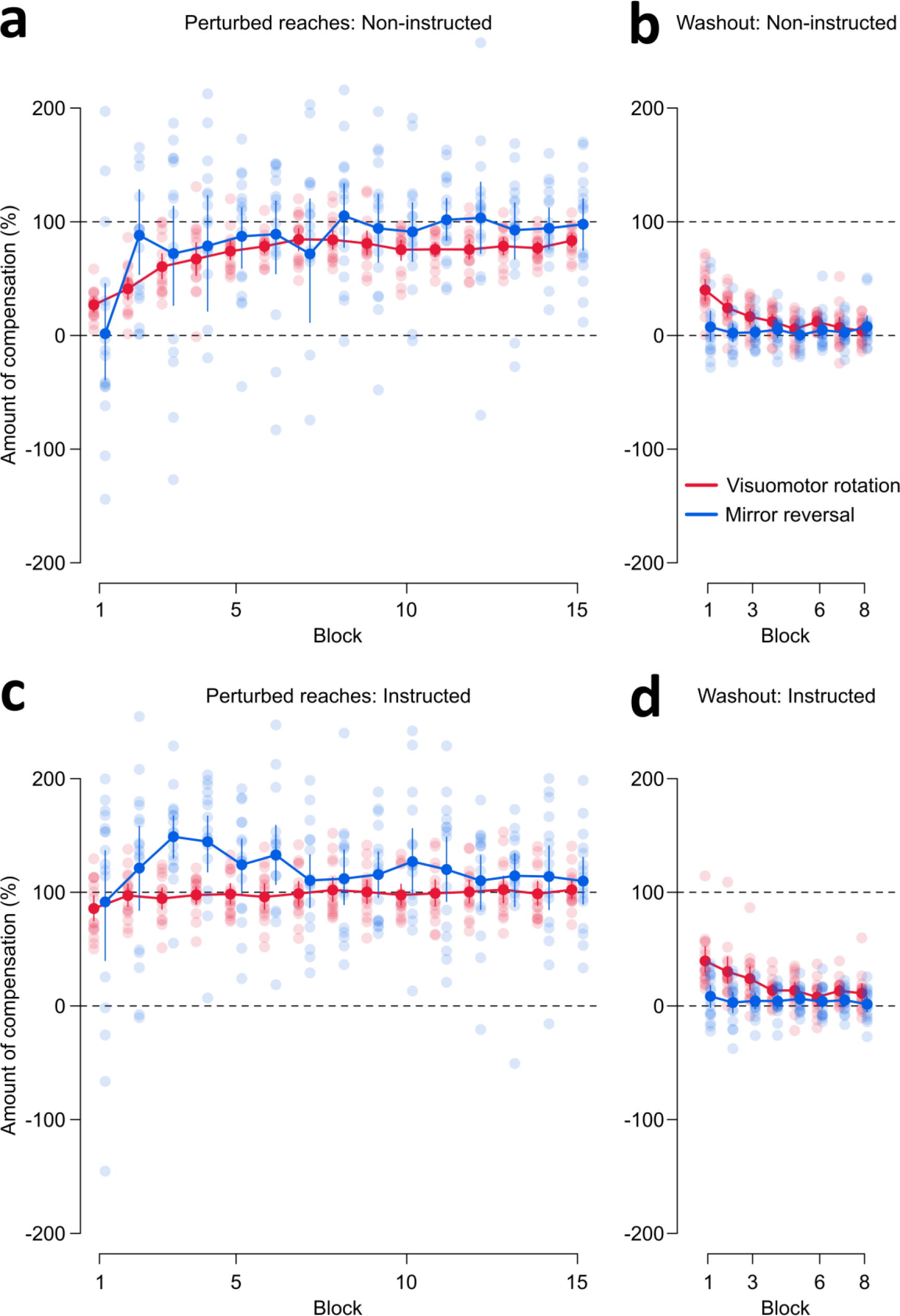
Reach performance during perturbed and washout trials for (a-b) non-instructed and (c-d) instructed participants. To make the two perturbations comparable, we converted the angular reach deviations to percentage of compensation. The grey dashed line at 100% indicates fully and successfully countering for the perturbation, while the grey dashed line at 0% indicates reaching directly to the target or no compensation. Individual data are shown, with solid dots and error bars indicating means and 95% confidence intervals. **(a)** Non-instructed participants learn to compensate for both perturbations, but show more variability for the mirror reversal. (**b**) Washout trials following perturbation training show evidence of reach aftereffects and de-adaptation following rotation training only. (**c**) Receiving instructions and a strategy to compensate for the perturbation provides an immediate advantage in learning. However, inter-participant variability is still larger for the mirror reversal. **(d)** Aftereffects and de-adaptation for instructed participants are only observed following rotation training.

We then investigate reach aftereffects using the washout trials. As expected, we observe reach aftereffects and gradual de-adaptation following rotation training but not after training with the mirror reversal (Fig. 2B). Given the lack of aftereffects in mirror reversal washout, we confirm the presence of reach aftereffects with a paired t-test that compares the percentage of compensation during the last block of aligned reaches, with the first block of rotation or mirror reversal washout trials. We find a difference between aligned and rotation washout trials (BF > 4·10^4^), but no evidence for a difference with mirror reversal washout (BF = 1.277). In short, definitive evidence for reach aftereffects are only observed following rotation training.

For rotations, instructions about the nature of the perturbation and strategies to compensate for it provide an advantage in early learning ^[13,23,26]^. Here, we observe that instructions provide a similar advantage for mirror reversal learning, as participants immediately learn to compensate (Fig. 2C). However, variability in learning is still larger compared to rotated trials. We then test for reach aftereffects (Fig. 2D) using a paired t-test that compares the percentage of compensation during the last block of aligned reaches, with the first block of each perturbation’s washout trials. We find a difference between aligned reaches and rotation washout (BF > 4·10^3^), and no evidence for a difference with mirror reversal washout (BF = 1.905). Thus, we only find conclusive evidence for aftereffects following rotation training in instructed participants.

We also investigate learning using reaction time (RT), movement time (MT), and path length (PL). For these three dependent variables (Fig. 3), we conduct paired t-tests comparing the first and last block of perturbation training with the last block of aligned baseline trials. As expected, participants have slower RTs during the first block of training in the rotation perturbation (BF = 7.557), and have much slower RTs for the mirror reversal (BF > 10^4^). Given that initial mirror RTs are much slower than rotation RTs (Fig. 3A), we also find that only rotation RTs return to aligned baseline levels at the end of training (rotation: BF = 0.256; mirror reversal: BF = 3.027). For movement execution time (Fig. 3B), we observe faster MTs in the first and last blocks of rotation trials compared to aligned trials, while mirror reversed trials show a difference with aligned MTs during the last block of training but not during the first training block (BF = 0.259). However, MTs are overall comparable across trial types (Fig. 3B). For path length, we observe greater inter-participant variability in mirror reversal trials (Fig. 3C) but find no effects for both perturbations. Taken together, movement initiation is generally slower for mirror reversed trials.

**Figure 3.**
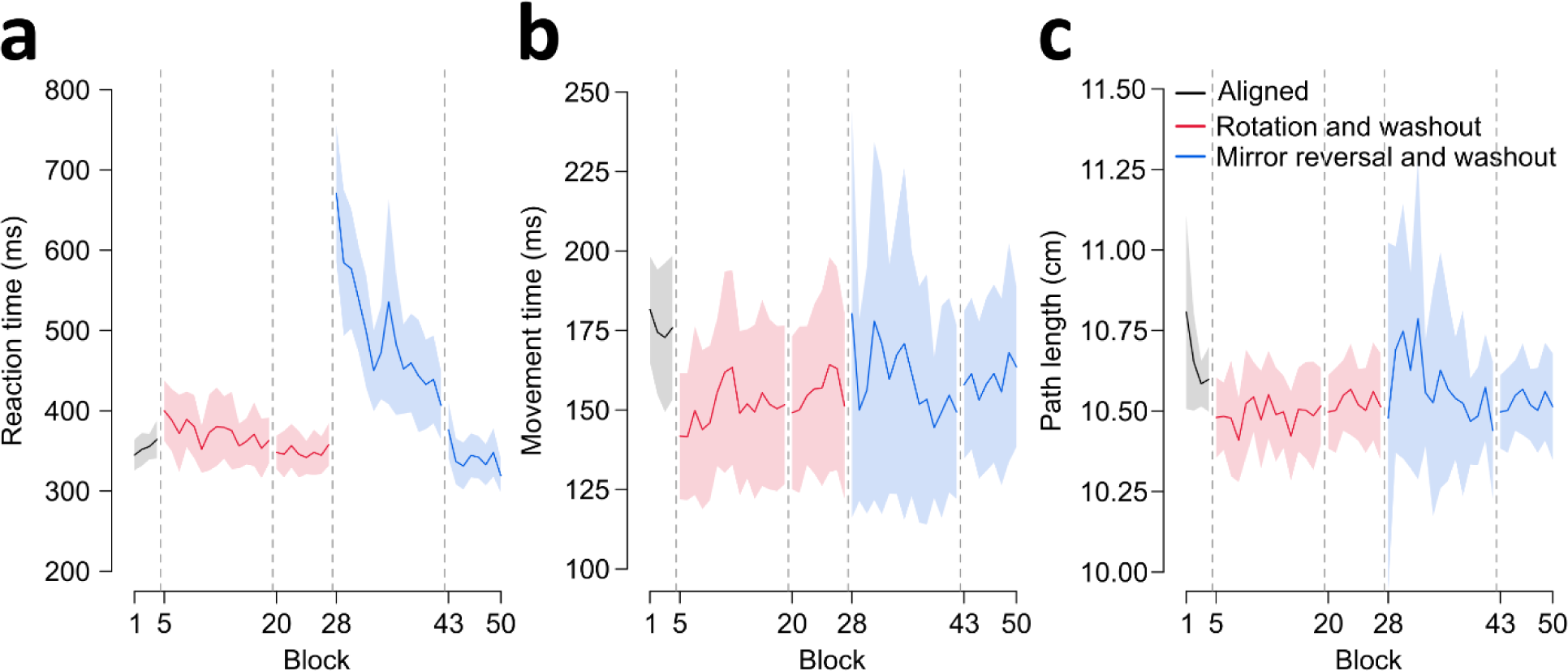
Movement measures during aligned, perturbed, and washout trials. (**a**) Reaction time (RT) refers to the time elapsed between the go signal onset and when the hand-cursor has moved 0.5 cm away from the start position. RTs across the different trial types are shown. Participants show slower RTs for perturbed trials, but only rotation trial RTs return to aligned baseline levels. RTs in the mirror reversal are much slower compared to rotation RTs. (**b**) Movement time (MT) is the time elapsed between the first sample when the hand-cursor is >0.5 cm away from the start position and the first sample when the hand-cursor is greater than the start-to-target distance. MTs for the rotation trials are faster than aligned baseline levels. Mirror MTs only show a difference with aligned MTs during the last block of training. (**c**) Path length (PL) refers to the total distance, given x and y coordinates of all recorded points on the reach trajectory, travelled between movement onset and offset. The shortest path to the target is a straight line spanning the start-to-target distance (9 cm). Inter-participant variability is larger for path lengths during the mirror reversal perturbation, but overall there are no differences in path length across trial types. For (**a-c**), each block shows the average of six trials. Solid lines and shaded regions are means and 95% confidence intervals across participants.

### Movement measures show quick learning in an online mirror reversal task

We investigate de novo learning further and incorporate the mirror reversal perturbation in an online experiment (Fig. 1E-1F). Participants control a cursor with their mouse or trackpad to reach towards targets while using their dominant or non-dominant hand. Since we only investigate the mirror reversal here, we are not constrained to use target locations that produce comparable perturbation magnitudes as in the rotation for our tablet experiment (around 30°; Fig. 1C). Furthermore, since we found that participants quickly compensated for the mirror reversal, with many of them over-compensating for the perturbation (Fig. 2), we want to investigate whether this quick learning is related to target location relative to the reversal axis. Thus, we placed a target 5° away from the axis (near target, Fig. 1F), a target at the other extreme and far from the reversal axis (175°, far target), and one along a perfect diagonal (45°, middle target).

Our statistical analyses include both target locations and blocks of trials as factors in different target X block within-subjects ANOVAs, where we compare learning on three dependent measures: percentage of compensation, completion time, and path length (Figs. 4-5). To keep our discussion of the results simple, we first describe block effects which address our primary focus on de novo learning, retention, and generalization. Then, we summarize results describing the impact of target location on learning.

**Figure 4.**
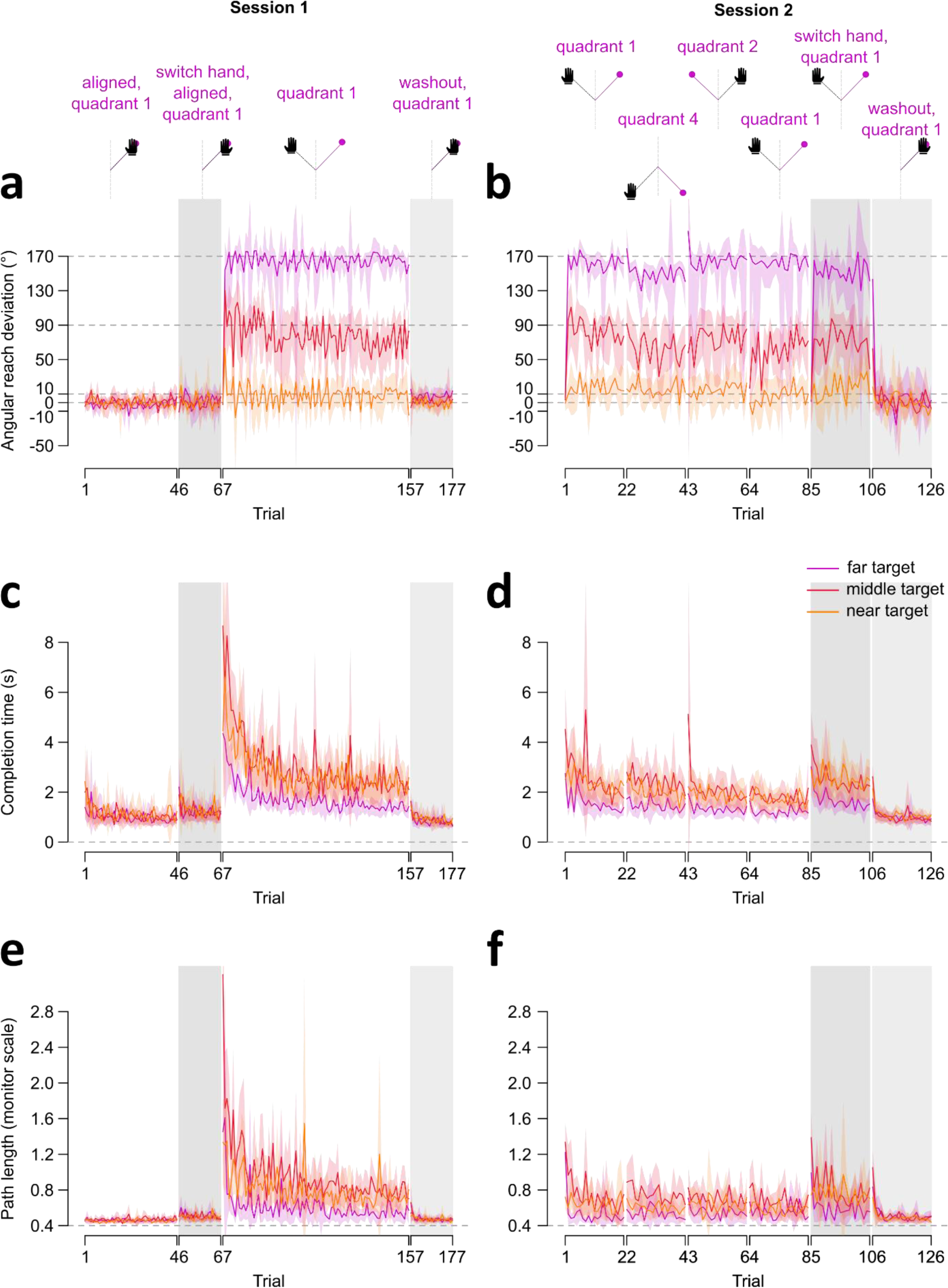
Reach performance, completion time, and path length across trial types in both sessions. Results from Session 1 are shown on the **left** (**a,c,e**) while those for Session 2 are shown on the **right** (**b,d,f**). On the top row, we depict the main task differences within and across sessions. **Left:** Session 1 began with 45 trials of aligned movements performed with the dominant hand, followed by 21 trials of aligned reaches using the non-dominant hand. Participants switch back to their dominant hand to complete 90 mirror reversed trials, followed by 21 washout trials. **Right:** Results of Session 2 to test learning retention from session 1. Session 2 began with 21 perturbed trials to targets in quadrant 1 (similar to the training quadrant in session 1). We then test for generalization with 21 perturbed trials to different target locations on the same (quadrant 4) or opposite (quadrant 2) sides of the mirror axis. To prevent decay in learning, we have them perform another set of 21 reaches to quadrant 1, before testing for intermanual transfer with their untrained or non-dominant hand (21 trials). The session ends with 21 washout trials using the non-dominant hand. Trial regions shaded in grey indicate washout trials and switching to the untrained hand in both sessions. **(a-b)** Angular deviation of the hand. Grey dashed lines at 10°, 90°, and 170° indicate the direction the hand must deviate to counter for the perturbation fully and successfully in near, middle, or far target locations respectively. Grey dashed lines at 0° indicate no compensation. (**c-d**) Completion time, reflects the time elapsed between target onset and target acquisition (RT plus MT). (**e-f**) Path length (PL). In the online study, PL reflects the total distance, given x and y coordinates of all recorded points on the cursor trajectory, travelled from the start position to the acquired target. Given that the distance between target and start position was 40% of the screen height (see methods), PL is normalized to be proportional to this minimum start-to-target distance (grey dashed line, at 0.4 height units). In all plots, solid lines and shaded regions represent means and corresponding 95% confidence intervals.

**Figure 5.**
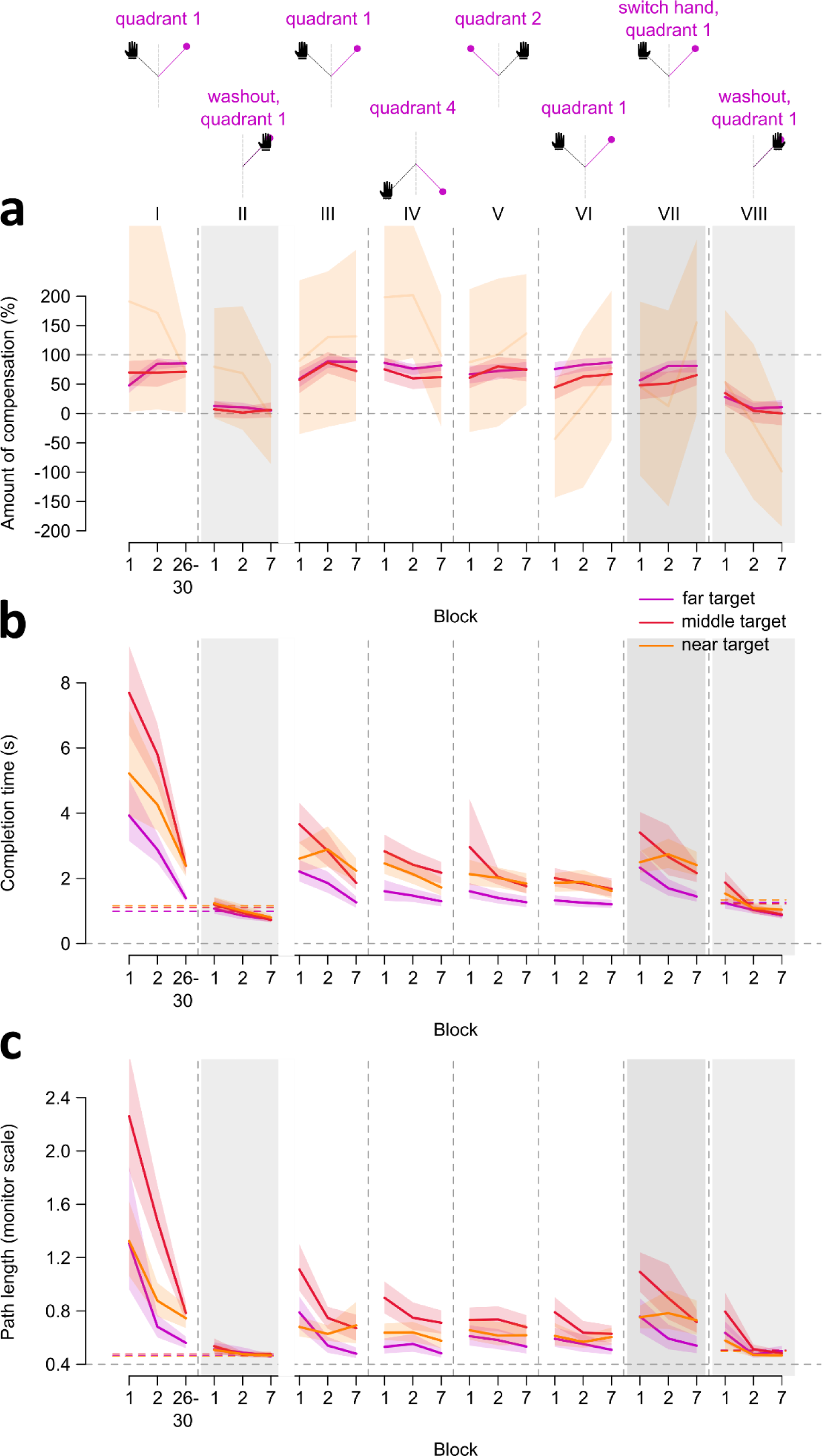
Percentage of compensation, completion time, and path length across blocks of trial types in both sessions. On the top row, we depict the main task differences within and across sessions, along with roman numerals to identify the specific trial blocks we compare in our analyses. The first set of blocks (column I) are the mirror reversed trials completed with the dominant hand in session 1, followed by the first, second, and last blocks of trials (column II) during session 1 washout. The remaining sets of blocks (columns III - VIII) are from session 2, where participants reached to targets in different quadrants, switched to using their non-dominant hand, and ended with washout trials using the non-dominant hand. Solid lines and shaded regions represent means and corresponding 95% confidence intervals. Regions of blocks shaded in grey indicate washout trials in both sessions and switching to the untrained hand in session 2. (**a**) Percentage or amount of cursor compensation, as described in Fig. 2, to make deviations to the far, middle, and near targets comparable, with 0% indicating no compensation and 100% perfect compensation (dashed lines). However, this normalization led to widely inflated variability for the near target, for reasons described in the methods. Thus, although we show data for the near target here (lighter shade of orange), we only report analyses comparing compensation between the far and middle targets. (**b-c**) Normalization is not required for completion time (**b**) and path length **(c)**. As such, we report comparisons across all three targets. Colored dashed lines indicate completion time and path length measures during baseline aligned trials for each target, while participants use either the dominant hand (columns I-II) or non-dominant hand (column VIII).

We first assess learning as participants train with the mirror reversed perturbation using their dominant hand. When normalizing the angular reach deviations (Fig. 4A) into percentages of compensation (Fig. 5A column I), we observe large inter-participant variability for near target percentages. We therefore exclude the near target from analyses comprising the percentage of compensation (discussed in Methods). As such, we only compare the far and middle targets across three different time points of mirror reversed training (Fig. 5A column I: blocks 1, 2, 26-30). However, we incorporate all three targets in analyses of completion time and path length, as normalization is not necessary for these measures (Fig. 5B-5C). For compensation, we surprisingly observe immediate compensation of the perturbation after one trial and consistent compensatory movements thereafter (Fig. 4A; inconclusive evidence of a block effect: BF_incl_ = 1.240). This is attributable to the lack of experimenter control in an online experiment, such that participants explore the workspace and figure out the perturbation. We find this behavior here and in a previous experiment version with over 600 participants ^[51]^. However, we observe a more typical, yet still fast, learning curve for completion time and path length for these mirror-reversed movements (Fig. 4C, 4E, 5B-5C). After just 90 mirror-reversed trials, we observe a notable improvement in completion times, initially exceeding 4 seconds, now reduced to approximately 2 seconds (Fig. 4C). Similarly, path length decreases from over 120% of screen height to approximately 60% of screen height (Fig. 4E). Although these measures do not fully return to baseline levels (aligned trials: mean completion time = 1.08 sec and path length = 47% of screen height), they are noticeably closer to baseline by the end of training (completion time: mean difference = 0.98 sec, BF > 2 ·10^18^, path length: mean difference = 23% of screen height, BF > 10^20^) compared to the beginning of training (completion time: mean difference = 4.60 sec, BF > 10^21^, path length: mean difference = 117% of screen height, BF > 5 ·10^14^). The initial values and amount of reduction in both measures differ across the three targets, with evidence for a target X block interaction for both completion time (BF_incl_ = 4.090) and path length (BF_incl_ = 20.690), which we discuss in a later section. Taken together, participants moved the cursor faster and more directly towards all targets as training progressed, suggesting that participants quickly learned to compensate for the mirror reversal.

We then examine washout trials following mirror reversed training with the dominant hand. To do so, we compare the first two blocks of washout trials (Fig. 5 column II) with aligned reaches. For compensation, we find no block effect (BF = 0.088), suggesting no aftereffects are present. For completion time and path length we find that movements during the initial block of washout are around 10% longer in time and distance than those in aligned trials, but these small differences disappeared by the second block of washout (strong evidence for a block effect in completion time: BF > 3 ·10^4^ and path length: BF = 831.79). These small and transient changes during washout are unlikely to be indicative of learning aftereffects. Consequently, we do not find definitive evidence supporting the existence of reach aftereffects following mirror reversal learning.

### De novo learning is retained across multiple days, shows intermanual transfer, and generalizes across the workspace

In the second session, we assess the retention of mirror reversed learning from the prior session, and evaluate the extent of generalization to the opposite hand and across different workspaces (Fig. 4 right side). However, the quick and near complete compensation observed in session 1 complicates the interpretation of similar compensation levels in session 2 (Fig. 4A-4B, 5A). This also applies to the interpretation of generalization across the hands or workspaces. Consequently, our primary focus will be on completion time and path length to examine both retention and generalization aspects.

For retention, we compare the first block of mirror reversed reaches in session 2 (Fig. 5 column III), with both the first and last training blocks from session 1 (Fig. 5 column I). We find both completion time and path length measures during session 2 are much smaller compared to initial learning in session 1, suggesting retention. That is, both measures in the first block of session 2 appear 50% shorter than those in the first block of session 1. This substantial retention, however, is not complete, as reaches in session 2 show initially longer measures than those at the end of session 1, before dropping to the same levels by the last block of that set of trials in session 2 (Fig. 5B-5C column III). These results are supported with strong evidence of a block effect (completion time: BF_incl_ > 10^25^, path length: BF_incl_ > 3 ·10^14^). Furthermore, these effects are modulated by target, with strong evidence for a target X block interaction (completion time: BF_incl_ = 10^4^, path length: BF_incl_ = 214.660). Taken together, measures of completion time and path length show evidence of partial learning retention even after three or more days.

We then examine whether learning while using the dominant hand transfers to the untrained and non-dominant hand. However, before we assess intermanual transfer, we first test for an effect of hand used in controlling the cursor during aligned baseline reaches (Fig. 4C,4E). We compare the dominant and non-dominant hands across different time points (first, second, and last blocks), and find strong evidence for a hand effect in both completion time (BF_incl_ > 10^3^) and path length (BF_incl_ > 4·10^12^). On average, completion time and path length were only 0.19 sec faster and 4% of screen height shorter for the dominant hand. (Fig. 4C,4E). This suggests that the dominant hand shows slight advantages for movement planning and execution. Given this small difference between hands, we remove this confound from our assessment of intermanual transfer by subtracting the mean completion time and path length measures from the first block of mirror reversed reaches using the untrained hand (Fig. 5 column VII). We then compare reaches in this first block of the untrained hand with both the first and last training blocks from session 1 (Fig. 5 column I). We find that both completion time and path length measures for the untrained hand are 50% shorter compared to initial learning in session 1, suggesting transfer of learning. These findings are confirmed with strong evidence for a block effect (completion time: BF_incl_ > 10^26^, path length: BF_incl_ > 2·10^14^). We also find strong evidence for a target X block interaction (completion time: BF > 2·10^4^; pathlength: BF = 196.430), and are illustrated with the means and confidence intervals displayed in figure 5B-5C. Despite this, both measures during the first block of reaches with the untrained hand are longer compared to the end of training in session 1. Thus, it seems that there is only partial transfer of learning across hands.

Our next step is to assess whether learning generalizes across the workspace. It is clear from figures 4D, 4F, and 5 columns IV-V, that initial completion time and path length measures in the novel workspaces are not only substantially lower compared to initial training in session 1 (Fig. 5 column I), but also seem to benefit from the first set of reaches in session 2 (Fig. 5 column III). We therefore conduct comparisons exclusively within session 2. Specifically, the initial set of reaches to quadrant 1 in the second session (column III) serve as the baseline and original trained workspace, against which we assess learning in the other novel quadrants. We compare the first and last blocks of the original training reaches with the first block of reaches to different test targets on either the same side of the mirror axis (column IV; Fig. 1E-1F), or opposite side of the axis (column V). For targets on the same side of the axis, we find that completion time and path length are overall lower for the test block compared to the first training block, confirmed with strong evidence for a block effect (completion time: BF_incl_ > 6·10^10^, path length: BF_incl_ > 2·10^7^). We also find a target X block interaction for both measures (completion time: BF_incl_ = 83.810; path length: BF_incl_ = 558.020). Thus, learning seems to generalize to targets on the same side of the mirror axis. For targets on the opposite side of the axis, completion time and path length measures show a similar pattern of results as in the previous set of targets, where we also find a block effect for completion time (BF_incl_ > 3·10^3^) and path length (BF_incl_ > 2·10^8^), as well as a target X block interaction for path length (BF_incl_ > 10^3^). Therefore, learning also generalizes to targets on the opposite side of the mirror axis.

Finally, participants complete washout trials to the trained targets but using the non-dominant hand. We compare the last block of non-dominant hand aligned trials in session 1 with washout blocks 1 and 2 using the non-dominant hand in session 2 (Fig. 5 column VIII). We find that the switch between perturbed to aligned reaches was not so smooth, such that the first washout trial with the non-dominant hand produced deviations consistent with the presence of aftereffects (Fig. 4B, 5A column VIII). However, by the second trial, there are no further deviations, suggesting that there are no aftereffects (Fig. 4B). Consequently, we find a block effect for compensation (BF_incl_ = 11.240), completion time (BF_incl_ > 3·10^13^), and path length (BF_incl_ > 2·10^14^). We also find strong evidence for a target X block interaction for both completion time (BF_incl_ = 321.310) and path length (BF_incl_ = 59.780). Deviations in the first trial are due to participants switching directly from perturbed to washout trials without any breaks. Overall, however, we do not find definitive evidence of aftereffects with the non-dominant hand.

### Movement measures depend on target location relative to the mirror axis

For the different statistical tests we have previously reported, we find that target location contributes to a significant interaction with the different blocks of trials considered. This is true for our comparisons during training in session 1 (Fig. 5 column I), for retention (column III), generalization across the workspace (columns IV-V), intermanual transfer (column VII), and washout in session 2 (column VIII). In all these tests, completion time and path length are overall lower for the far target (purple curve), compared to the middle (red) and near (orange) targets (Fig. 4C-4F, 5B-5C). That is, we observe that the purple curve falls below the confidence intervals of the other targets across most blocks. Thus, it seems that it is easier to compensate for the mirror perturbation when targets are placed far from the mirror axis.

## Discussion

We test different behavioral measures that distinguish motor adaptation from de novo learning. Particularly, we implement a within-subjects design where participants train with both a visuomotor rotation and mirror reversal perturbation, to investigate adaptation and de novo learning respectively. Participants’ learning for one perturbation does not affect the other, suggesting that the two motor learning types are independent. Mirror reversal learning is also more variable across participants, but develops as quickly and to a comparable asymptote as rotation adaptation. Furthermore, we find evidence for reach aftereffects following rotation training only, suggesting greater implicit contributions during adaptation. Consequently, explicit contributions are greater in de novo learning, as shown with slower movement initiation times. Despite an already fast learning rate for the mirror perturbation, instructions do lead to immediate and full compensation. Our larger sample from the browser-based study provides more evidence for de novo learning mechanisms. Although there were different error magnitudes from the three target locations, we find evidence that learning occurs quickly and does not produce reach aftereffects. Furthermore, this learning is partially retained across multiple days and partially transfers to the untrained hand and to targets across the workspace. Finally, movement measures show that it is easier to move towards targets when they are about perpendicular with the mirror axis, and harder when targets are placed along a diagonal. All these behavioral measures suggest that de novo learning has distinct underlying mechanisms compared to adaptation.

Adaptation results from an efferent-based component, where the predicted outcome of a motor command is updated to match the experienced visual outcome of the movement ^[2,4,19–20]^. Modifications to such an existing control policy typically progress within several trials to get back to previous levels of performance ^[1–4]^. Then once the perturbation is taken away, it also takes several trials to de-adapt. These are known as reach aftereffects, and is typically considered as evidence for an implicit contribution to learning ^[1,12–13,25–26]^. However, in de novo learning, adaptation based on sensory prediction errors is counterintuitive, as one needs to increase the hand-cursor discrepancy to reach towards a target ^[5,10–11,29,31–34]^. Instead, a different and new control policy must be established ^[5,33]^, and one can simply switch between policies depending on the presence of the perturbation. If different mechanisms underlie learning in the two perturbation types, then learning in one perturbation should not affect the other ^[52]^. Further, we do not expect aftereffects following de novo learning. In the tablet experiment, we have participants train in both perturbations. We find no perturbation order effects, suggesting that different learning mechanisms operate for each perturbation. Moreover, in both our studies, we find no definitive evidence of reach aftereffects following mirror reversed training. Although our results from both studies may seem indicative of, or show weak evidence for, aftereffects following mirror reversed training, these are smaller compared to those we observe following rotation training and are transient such that we do not observe gradual de-adaptation across washout. Notably, we do not find aftereffects regardless of how large the visual error is, based on the target distance from the mirror axis. Taken together, adaptation is independent from de novo learning, and implicit contributions to learning are efficient for adaptation but counterproductive for de novo learning.

Explicit components contribute more for de novo learning. In adaptation, larger explicit contributions to learning are observed with larger perturbations and strategies to compensate for the perturbation ^[23–26]^. We do not compare the relative contributions of explicit and implicit learning, but we do take the absence of implicit aftereffects as evidence of mostly explicit learning for the mirror perturbation. Moreover, implicit components are usually limited in their contribution to adaptation ^[4,24,53–54]^ with explicit components contributing to full compensation. As we find comparable asymptotic learning for the mirror reversal and rotation perturbations in the tablet study, it is likely that explicit components contribute more towards full compensation. We also find that providing a strategy to counter the perturbation leads to immediate compensation for the mirror reversal. For a more direct measure of explicit contributions, we measure reaction times for the reaching movement. We expect that more preparation for the upcoming movement will lead to longer reaction times and less errors ^[5,11]^. In our tablet study, we find that reaction times are much slower for the mirror reversal compared with the rotation. Moreover, reaction times for the mirror reversal do not go back to baseline levels by the end of training. These findings suggest that participants take more time to prepare the upcoming movement. Additionally, we measure path length and find that participants show more variability in their paths for the mirror reversal early in learning, suggesting that participants are figuring out the optimal path towards the target. Such measures provide insight into how much more cognitively demanding it is to compensate for the mirror reversal.

Cognitively demanding tasks, like the mirror reversal, require more practice. Previous experiments have used hundreds of trials and multiple days to show how long mirror reversed learning develops ^[5,11,27–28,30]^. In the tablet study, we show full compensation within 90 trials. However, variability is large across both non-instructed and instructed participants. This suggests that even though participants figure out a global strategy to compensate for the mirror reversal, more practice is required to make reaching directions more precise and consistent across trials. For the online study, we observed almost immediate learning after one trial. Upon closer inspection, and with data from more than 600 people ^[51]^, we find that this fast learning occurs for people who explore the workspace to figure out the perturbation. However, such immediate learning is not observed in online adaptation studies ^[14,55]^, suggesting that exploration is unique for de novo learning. Finally, the effect of cognitive demand on movement seems to be target-dependent. The faster completion times and shorter path lengths for the far target compared to the middle target suggest that it is easier to compensate for the mirror reversal when simply flipping along the left-right direction, rather than combining this with a diagonal movement. Thus, the amount of practice, exploration, and target location are all factors that may affect reaching performance in a mirror reversal task.

Retention is also expected to be different between adaptation and de novo learning. Adaptation typically decays with the passage of time ^[36]^ and with washout trials ^[4,39]^. Further, even if adaptation may be retained for a few days or up to a year ^[4,37–38]^, retention is only partial. Adaptation is also characterized with savings, where people commit large initial errors upon experiencing the perturbation a second time, but show faster re-learning rates ^[40–41]^. Conversely, de novo learning is characterized with offline gains, where the starting performance is similar or better than the end of the previous session ^[5–7,10–11]^. In the online experiment, we observe quick and near complete compensation in session 1, thereby complicating interpretation of similar compensation levels we observe in session 2. We instead find that completion time and path length are substantially shorter at the start of session 2 compared to session 1 start, showing that learning is retained across three or more days, even though participants performed washout trials at the end of session 1. Moreover, both measures at the start of session 2 are longer than those at the end of training, suggesting that retention is partial. However, we observe that these measures immediately return to similar levels as in the end of session 1 in the following trials. Taken together, although we do not find conclusive evidence for offline gains, our findings still show that de novo learning is fairly persistent.

Learning also generalizes differently between de novo learning and adaptation. In adaptation, narrow generalization patterns are observed with aftereffects ^[1,42–43,56–57]^. There are no aftereffects to compare generalization patterns in the mirror reversal. However, generalization patterns are also expected to be narrow, since implicit learning is target-dependent ^[10,29]^. In the current study, we tested participants on corresponding target locations (i.e., far, middle, near) on either side of the mirror axis. Despite targets being located on novel workspaces, we find that completion time and path length are overall shorter than those in the initial training quadrant, suggesting transfer of learning. While we cannot rule out implicit contributions, it is likely that explicit components contribute to this transfer. Another form of generalization, intermanual transfer, is asymmetric for adaptation ^[44–48]^, likely due to error attribution for the more unreliable non-dominant hand ^[13,58–59]^. Asymmetric intermanual transfer is not observed for tasks with reversed visual feedback ^[49–50]^, but these studies reversed feedback of the actual hand, which is different from our study’s mirror reversal. In our study, we investigate intermanual transfer from the dominant and trained hand to the non-dominant or untrained hand. We find slight advantages for movement planning and execution for the dominant hand. We thus account for this difference and find that completion time and path length measures for the non-dominant hand are overall shorter than initial training with the dominant hand. However, this transfer is only partial as movement measures are longer than the end of training with the dominant hand. Thus, mirror reversal learning transfers across the workspace and hands.

In summary, we show learning, reach aftereffects, retention, and generalization patterns for de novo learning which differ from patterns usually observed with adaptation. Particularly, de novo learning is more variable, has more explicit contributions, and produces different movement initiation and execution times depending on target locations. Retention for de novo learning is more robust, and learning transfers across the workspace and to the untrained hand. These behavioral differences are likely due to distinct neural mechanisms underlying each motor learning type. Adaptation, for example, is dependent on the cerebellum ^[2–3,22,60–61]^, while de novo learning seems to depend on areas like the basal ganglia ^[9,62]^. Future research should probe the different neural processes underlying de novo learning, to better understand how efficient control policies are established. Overall, multiple behavioral measures provide insight into how new motor skills are acquired and how well-known motor skills are adapted.

## Methods

### Participants

32 healthy adults (25 female, 3 left-handed, *M*_Age_ = 20.0, *SD*_Age_ = 3.03) participated in the tablet experiment (study 1), but only half of the participants received instructions about the nature of the perturbation and a strategy to counter for it (instructed: *n* = 16, 12 female; non-instructed: *n* = 16, 13 female). For the online experiment (study 2), 63 healthy adults (44 female, 1 identified as neither sex, 5 left-handed, 1 identified as ambidextrous, *M*_Age_ = 22.0, *SD*_Age_ = 5.79) participated in session 1, and 48 participants (36 female) returned for session 2. While the sample size in study 1 is typical for laboratory studies ^[5–6,10–11,33]^, we took advantage of the ability to add more participants for study 2 as it was a browser-based experiment. All participants gave written informed consent prior to participating. All procedures were in accordance with institutional and international guidelines, and were approved by York University’s Human Participants Review Committee.

## Study 1: Tablet experimental set-up

### Apparatus

Participants sat on a height-adjustable chair in front of a digitizing tablet (Wacom Intuos3, 12” x 12” surface, resolution resampled by 1680 x 1050 pixels at 60 Hz) and a vertically-mounted monitor (Dell Technologies, 22” P2217 LED screen). The monitor (Fig. 1A) was located 55 cm from the tablet. An opaque shield was positioned on top of the tablet to occlude visual feedback of the participants’ arm. Participants used their right hand to move a digitizing pen across the tablet surface. A circular plastic stencil (20 cm in diameter, ^[63]^) was placed on top of the tablet surface and acted as a physical boundary for the arm movements.

### Stimuli

A cursor (white circle, 1 cm in diameter) represented the position of the digitizing pen on the monitor. Participants made ballistic reaching movements from the start position (grey circle, 1 cm in diameter at the centre of the screen) and had to slice through a target (blue circle, 1 cm in diameter) until they hit the circular stencil (Fig. 1). Targets were arranged radially, 9 cm from the start position. Target locations were dependent on participant order, perturbation order, and axis orientation. Each participant trained with both rotation and mirror perturbation types, where participants with even numbered IDs experienced the rotation before the mirror task and vice-versa for participants with odd numbered IDs. Each perturbation type corresponded to either the vertical or horizontal midline axis. As such, half the participants trained with the rotation perturbation on the horizontal axis, while the other half experienced rotation training on the vertical axis. Each axis and perturbation type had six corresponding training targets, either 7.5°, 15°, or 22.5° away from either end of the axis on its positive or negative side. This produced eight possible conditions to counterbalance across participants, perturbation order, and axis orientation (Fig. 1C). Targets were presented once in a shuffled order before being presented again, such that reach directions were evenly distributed across locations in all trial types. Trial types proceeded in a particular order across participants: familiarization, aligned reaches, perturbed reaches, and washout (Fig. 1B). Perturbed reaches and washout trials were repeated for the second perturbation type.

### Trial types

#### Familiarization

Participants kept the hand-cursor at the start position for 300 ms (Fig. 1D). The target location was then visually cued with a grey circle (1 cm in diameter), and participants had to hold their position for one second. After the hold period, the target cue turned blue, acting as the go signal to start moving. With the hand-cursor feedback shown continuously, participants then moved and sliced through the target. Once they hit the stencil, they were instructed to hold their reach endpoint position for one second. To ensure that participants performed ballistic movements, they had to reach through the target distance within 400 – 700 ms after the go signal onset. Additionally, they had to keep the hand-cursor within 1 cm of their reach endpoint. If they satisfied both conditions, the target disappeared, and the start position was presented as a blue circle. Otherwise, they heard a loud beep and the start position turned red. In either case, they then moved back to the start position to end the current trial. For familiarization trials, participants may only proceed to the next trial if they moved within the speed criteria and held their position at reach endpoint. One block of familiarization trials consisted of reaching to 12 target locations, which are the same targets they will experience in both the rotation and mirror perturbation types. Participants completed two blocks of familiarization trials, for a total of 24 trials.

#### Aligned reaches

These proceeded similarly as the familiarization trials. However, participants may continue to the next trial even if they did not satisfy the conditions for movement speed or position hold at reach endpoint. The start position still turned either blue or red at the end of the trial, to remind participants that they should move within the speed criteria and hold their reach endpoint position. One block of aligned reaches consisted of 12 target locations. Participants completed four blocks of aligned reaches, for a total of 48 trials. These trials served as baseline data for the perturbed reaches.

#### Perturbed reaches

Before these trials began, we ensured that the instructed participants understood the nature or the perturbation and how to counter for it, while the non-instructed participants were simply informed that the cursor would move differently and that they had to compensate for it. We perturbed visual feedback of the hand-cursor in two ways. In rotation trials, the cursor feedback was rotated 30° CW or CCW (counterbalanced across participants) relative to the hand position. To correct for this perturbation, participants had to move 30° in the opposite direction of the rotation. In mirror reversed trials, the cursor feedback was in the flipped direction of the hand position, relative to a mirror axis placed on either the vertical or horizontal midline. Correcting for this perturbation required moving towards the opposite side of the mirror axis. In both perturbation conditions, perfect compensation should result in the same set of movement dynamics. Participants trained with one perturbation type before training with the other. In each perturbation type, one block of trials consisted of six target locations. To ensure saturation of learning, participants completed 15 blocks of perturbed reaches, for a total of 90 trials. After training with each perturbation type, participants completed washout trials.

#### Washout trials

These proceeded similarly as the aligned reaches. However, one block of trials consisted of the same six target locations as in the corresponding training trials for each perturbation type. Participants completed eight blocks of washout trials, for a total of 48 trials. These trials are where we measured for reach aftereffects, or the persistent deviation of hand movements once the perturbation is turned off.

## Study 2: Online experimental set-up

### Apparatus

Study 2 was a browser-based experiment and participants used either their personal computer or laptop to complete the study. Throughout the study, participants use either their mouse (session 1: *N* = 19, session 2: *N* = 9) or trackpad (session 1: *N* = 44, session 2: *N* = 39) to control the cursor. Participants accessed the study using individualized anonymous links through York University’s Undergraduate Research Participant Pool SONA System. They completed an online questionnaire hosted on Qualtrics, where we collected information such as their screen resolution, demographic information, and included questions to check for which hand they used in the different experimental blocks. The questionnaire included embedded links to the experimental tasks, programmed using PsychoPy (version 2021.1.4) and hosted on the accompanying experiment server Pavlovia. Participants’ devices had a refresh rate of 60 Hz, which is standard for most laptop and desktop monitors. The experimental tasks were displayed on the web browser window in full screen mode.

### Stimuli

The size and position of the stimuli were dependent on each participant’s screen resolution and were scaled accordingly. Stimuli positions followed a Cartesian coordinate system, with the origin (0, 0) positioned at the centre of the screen. Negative values represented stimuli positioned down/ left relative to the origin, while positive values represented stimuli positioned up/ right. We scaled the stimuli relative to the participant’s screen height using the “height units” (h.u.) in PsychoPy. This meant that the upper and lower edges of the screen were always equal to ±0.5 h.u. (or 50% of screen height).

A cursor (black circle, 0.025 h.u. in diameter) represented the mouse or trackpad position on the monitor. Participants performed out-and-back reaching movements from the start position (white circle, 0.05 h.u. in diameter) located at the origin, to a target (white circle, 0.05 h.u. in diameter). Targets were arranged radially, located 0.4 h.u. or 40% of screen height from the origin. In session 1 (Fig. 1E-1F), there were three possible target locations located in quadrant 1 of a Cartesian coordinate system (5°, 45°, or 85° in polar coordinates). In session 2, we tested for retention and generalization across the workspace or hands. As such, the three possible target locations were positioned in either quadrant 1 (Fig. 1F; 5°, 45°, or 85° in polar coordinates), quadrant 2 (95°, 135°, or 175° in polar coordinates), or quadrant 4 (275°, 315°, or 355° in polar coordinates). Each of the three possible targets in a quadrant were presented once in a shuffled order before being presented again, to ensure evenly distributed reaches across trial types.

### Trial types

#### Aligned reaches

To begin each trial, participants had to move their cursor to the start position. Once the centre of the cursor was less than 0.05 h.u. away from the start position centre, a target appeared. Participants then moved towards the target until they acquired it, i.e., when the cursor centre was less than 0.05 h.u. away from target centre. They then moved the cursor back to the start position to continue with the next trial. Session 1 started with two sets of aligned reaches. In the first set, we instructed participants to control the cursor with their dominant hand, and reach to the three target locations in quadrant 1. Participants completed 15 blocks of aligned reaches with the dominant hand, for a total of 45 trials. In the second set, we instructed them to switch to using their non-dominant hand to control the cursor, while performing reaches to the same target locations. Participants completed seven blocks of aligned reaches with the non-dominant hand, for a total of 21 trials (Fig. 1E). Both sets of aligned reaches served as baseline data for the perturbed reaches.

#### Perturbed reaches

All perturbed reaches consisted of mirror reversed trials, where the cursor feedback was flipped in the left-right direction (mirror axis placed on vertical midline). In session 1, one block of trials consisted of the same three target locations as in the aligned reaches. Participants completed 30 blocks of perturbed reaches, for a total of 90 trials, using their dominant hand. In session 2, we measured for retention and generalization. As such, all sets of trials involved perturbed reaches (Fig. 1E). Participants returned after a minimum of three days (days apart: *M* = 4.77, *SD* = 2.52), and immediately performed 21 trials of perturbed reaches to quadrant 1 targets. They then completed 21 trials of perturbed reaches to quadrant 4 targets followed by 21 trials to quadrant 2 targets. To prevent any possible decay in learning, we had them perform another 21 trials of perturbed reaches to quadrant 1 targets. Finally, we instructed them to switch to their non-dominant hand, and perform 21 trials of perturbed reaches to quadrant 1 targets. Both sessions 1 and 2 ended with participants completing a set of washout trials.

#### Washout trials

These proceeded similarly as the aligned reaches, and were used to measure for any reach aftereffects. In each of the two sessions, participants completed seven blocks of washout trials to targets located in quadrant 1, for a total of 21 trials. However, participants used their dominant hand to control the cursor in session 1, and used their non-dominant hand to control the cursor in session 2.

## Data analysis

We used the different trial types to compare learning between the rotation and mirror perturbations in the tablet study, and to quantify retention and generalization of learning for the mirror perturbation in the online study. We report Bayesian statistics to show evidence in support of the alternative hypothesis over the null hypothesis (i.e., BF_10_ values). We include Bayes Factors (BF) for paired comparisons and inclusion Bayes Factors (BF_incl_) when assessing particular effects of predictors. We also report more detailed results from frequentist tests in our accompanying R notebook ^[51]^. All data preprocessing and analyses were conducted in R version 4.2.2 ^[64]^.

### Amount of compensation

We quantified reach performance using the angular reach deviation of the hand, which is the angular difference between a straight line connecting the start position to the target and a straight line connecting the start position to the point of maximum velocity in the reach. For the online study, we used a proxy for the point of maximum velocity, defined as the first sample after the cursor crosses 20% of the start-to-target distance. Regardless, the angular deviations for aligned reaches are expected to be close to zero. For both perturbed and washout reaches, we first corrected for individual baseline biases by calculating the average angular deviation for each target within each participant during aligned reaches, and subtracting this from angular deviations during perturbed or washout reaches. In the tablet study, the rotation was at a fixed magnitude of 30° while the magnitude of the mirror perturbation was dependent on how far the target was from the mirror axis. Therefore, although full compensation for the rotation was an angular reach deviation of 30°, full compensation for the mirror could either be 15°, 30°, or 45°. Consequently, in the online study, the different target locations corresponded to full compensation values of 10°, 90°, or 170°. Thus, to make the comparisons in perturbed reaches comparable, we converted the angular reach deviation measures during aligned, perturbed, and washout reaches into percentages.

#### Rate of change and asymptote of learning

With the percentage measures, we estimated the rate of change during perturbed reaches using an exponential decay function with an asymptote. The function is expressed as:

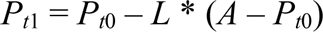

where the process value on the next trial (P_t1_) is equal to the current trial’s process value (P_t0_) minus the product of the rate of change (L) and error on the current trial (difference between asymptote, A, and the current trial’s process value). We constrained the rate of change (L) parameter to range [0, 1], and the asymptote (A) parameter to range [−1,2·max(data)]. We bootstrapped both parameters (1000 samples per fit) across participants to obtain a 95% confidence interval for each parameter.

For the tablet study, we fit the exponential decay function to the reach data of only the non-instructed participants during rotated and mirror reversed trials. Since inter-participant variability in mirror reversed trials was large, with some participants showing immediate learning after a given trial, we also attempted to fit the data using a step or logistic function. However, model comparisons showed that the exponential decay function provided the best fit for these perturbed trials (see R notebook ^[51]^). Thus, we compared the rate of change and asymptote parameters from the exponential decay function between the two perturbation types to quantify differences in learning. For the instructed participants in the tablet study, we were only interested in seeing the advantage of instructions on reaching performance. Given that the instructed participants did show an advantage in initial learning for both perturbations (Fig. 2C), we did not further analyze their perturbed reaches.

For the online study, it is inappropriate to fit the exponential decay function to the percentages of compensation, given that we observed immediate and consistent compensatory movements after one trial. We therefore compared compensation among target locations during the first, second, and last blocks of trials during learning (Fig. 5A). Using data from all three targets, we found no compensation effects across targets and blocks. However, the absence of effects may be due to the large inter-participant variability that occurs when reach deviations are converted into percentages, as shown in figure 5. For the near target, conversion into percentages uses a very small denominator, such that deviations of only a few degrees on either side of the target would correspond to a wide range of compensation values. Although the increased inter-participant variability in percentages does show some improvement with training (Fig. 5A column I), it still poses a challenge in discerning whether the compensation for the near target significantly diverged from that of the other two targets, and whether it changes with training. For this reason, we display the percentage of compensation data for the near target but omit it from analyses comparing compensation across targets that we report in this paper. Furthermore, this quick learning we observe in session 1 confounds our interpretation of similar compensation levels observed in session 2. As such, we focus on other descriptors of reaches for session 2 analyses (e.g., completion time and path length), as discussed below.

#### Reach aftereffects

For the tablet study, fitting the exponential decay function is inappropriate for mirror washout trials, since angular reach deviations are near zero and we do not observe de-adaptation. We therefore confirmed the presence of reach aftereffects by comparing the percentages of compensation during the last block of aligned reaches with the first block of each of the corresponding washout trials after rotated and mirror reversed training. We repeated the same set of analyses for the instructed participants. For the online study, we compared compensation in the first two blocks of washout trials with aligned reaches. Washout trials in session 1 were compared with aligned trials using the dominant hand, while washout trials in session 2 were compared with aligned trials using the non-dominant hand.

### Movement analyses

We investigated other measures of learning associated with the reaching movement, including reaction time, movement time, completion time, and path length.

#### Reaction time

In the tablet study, we defined reaction time (RT) as the time elapsed between the go signal onset and when the hand-cursor has moved 0.5 cm away from the start position. We compared RTs between the last block of aligned trials with the first and last blocks of rotation and mirror reversed training.

#### Movement time

In the tablet study, we defined movement time (MT) as the time elapsed between the first sample when the hand-cursor is >0.5 cm away from the start position and the first sample when the hand-cursor is greater than the start-to-target distance. We compared MTs between aligned and perturbed trial types, similar to how we analyzed the RTs.

#### Completion time

In the online study, there was no pause in between trials. One trial would end as participants moved close enough to the home position, and the target for the next trial would immediately appear. However, the return movement from that previous trial often overshot the home position, such that it appeared participants responded to the appearance of the target immediately. This made the definitions of reaction time used for the tablet study uninformative, and consequently that of movement time, as well as other ways to define RT and MT. We instead used completion time, defined as the time elapsed between target onset and target acquisition (RT plus MT). We compared completion time across different training blocks or trial types. For these analyses, we incorporated data from all three targets, as normalization is not necessary for this measure.

#### Path length

We defined path length (PL) as the total distance, given x and y coordinates of all recorded points on the reach trajectory, travelled between movement onset and offset. Movement onset and offset are the start and end movement times for the tablet study, and start and end completion times for the online study. The shortest path to the target is a straight line spanning the start-to-target distance. In the tablet study, we tested how PLs differed between the last block of aligned trials and the first and last blocks of perturbed reaches. In the online study, we compared PLs across different training blocks or trial types and incorporated data from all three targets.

### Data availability

Data and analyses scripts are available on Open Science Framework ^[51]^.

## Acknowledgements

This work was supported by NSERC for D.Y.P.H.; NSERC, OGS, and VISTA for R.Q.G. The funders had no role in study design, data collection and analysis, decision to publish, or preparation of the manuscript.

## Author Contributions

B.M.tH. and D.Y.P.H. designed the research. R.Q.G. collected the data. B.M.tH. and R.Q.G. contributed experimental and analytic code. R.Q.G. and B.M.tH. analyzed the data. R.Q.G. wrote the manuscript, which was carefully edited by all authors. The final version of the manuscript has been approved by all authors who agree to be accountable for all aspects of the work in ensuring that questions related to the accuracy or integrity of any part of the work are appropriately investigated and resolved.

## Conflict of Interest

The authors declare no competing interests.

## Notes

https://osf.io/786nf/

### Competing Interest Statement

The authors have declared no competing interest.

